## Supplementary figures and images for "PPanGGOLiN: depicting microbial diversity via a partitioned pangenome graph"

### S1 Fig

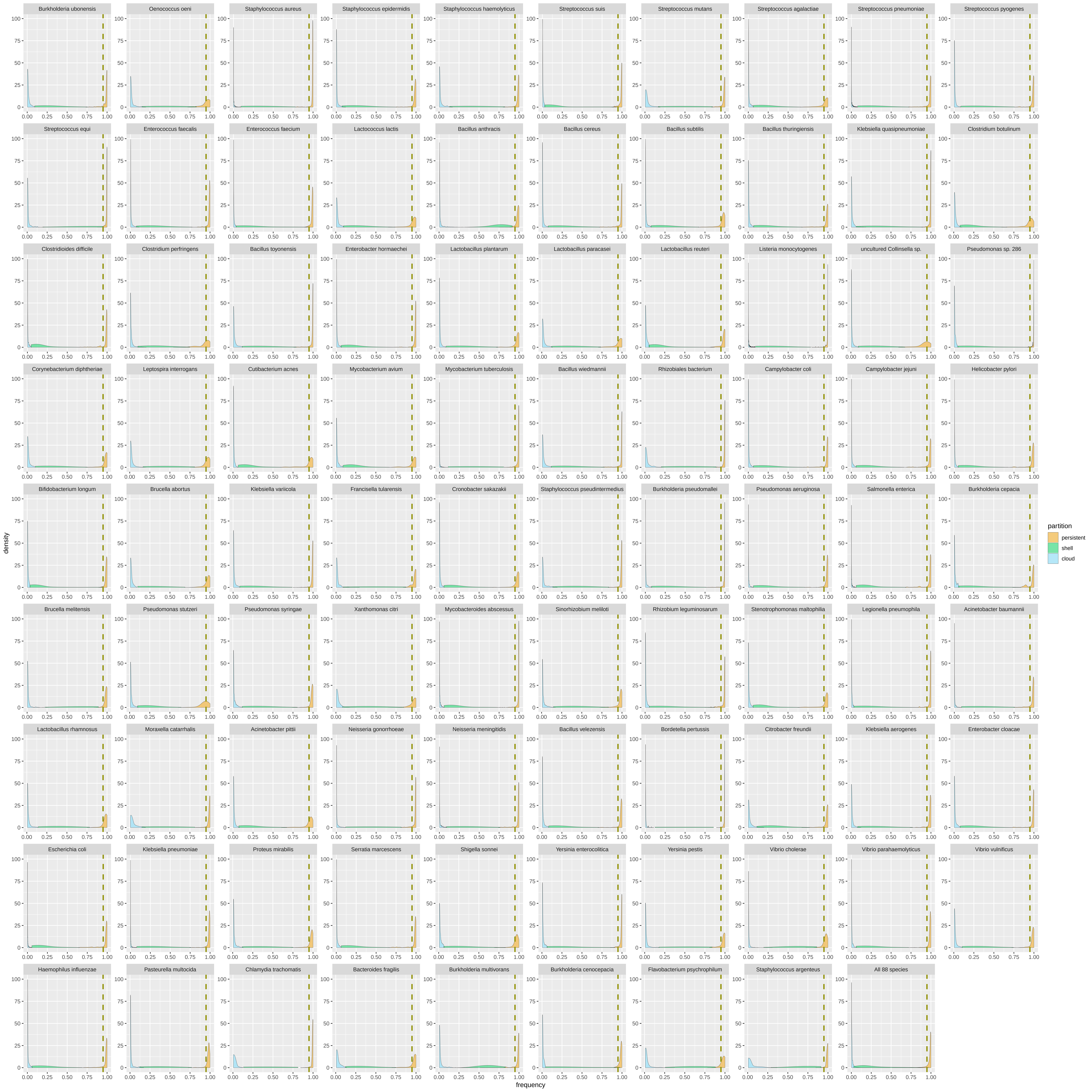

### S2 Fig

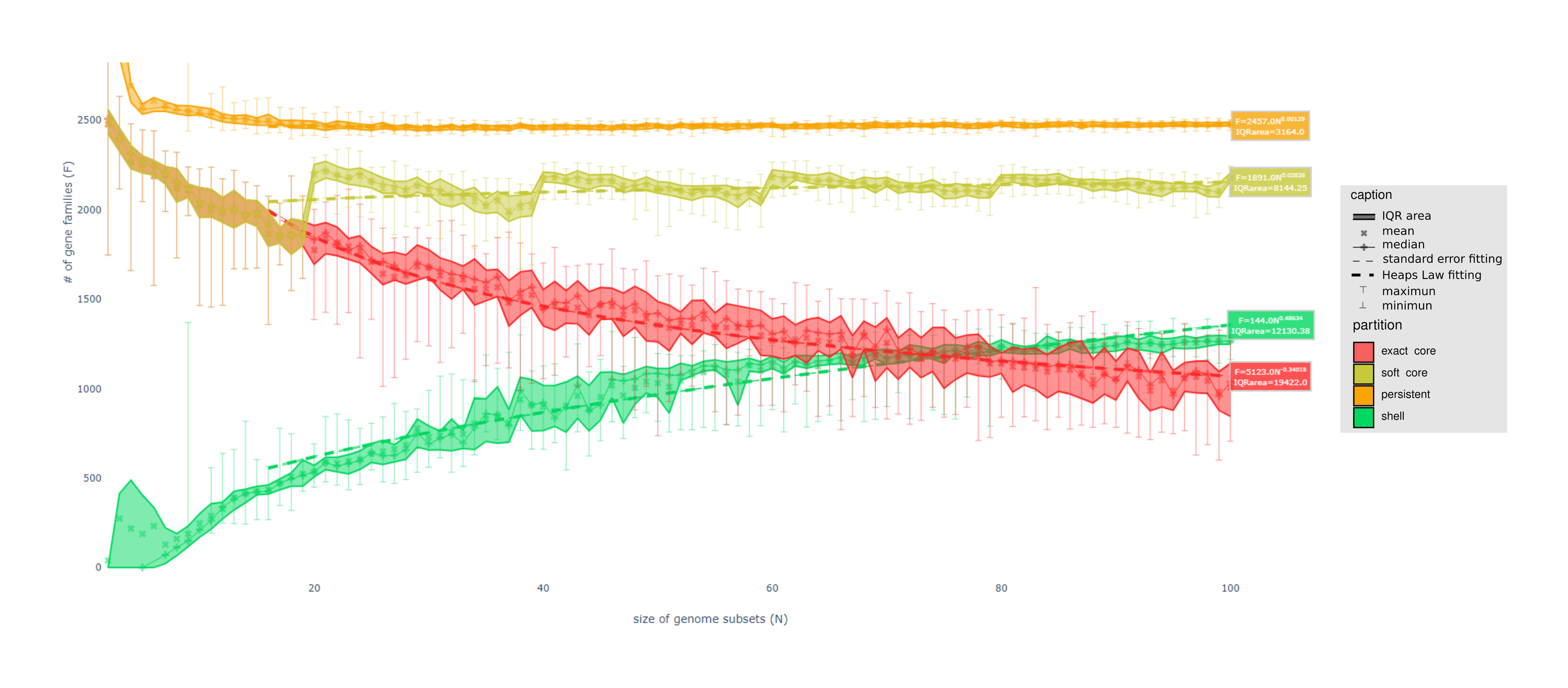

### S3 Fig

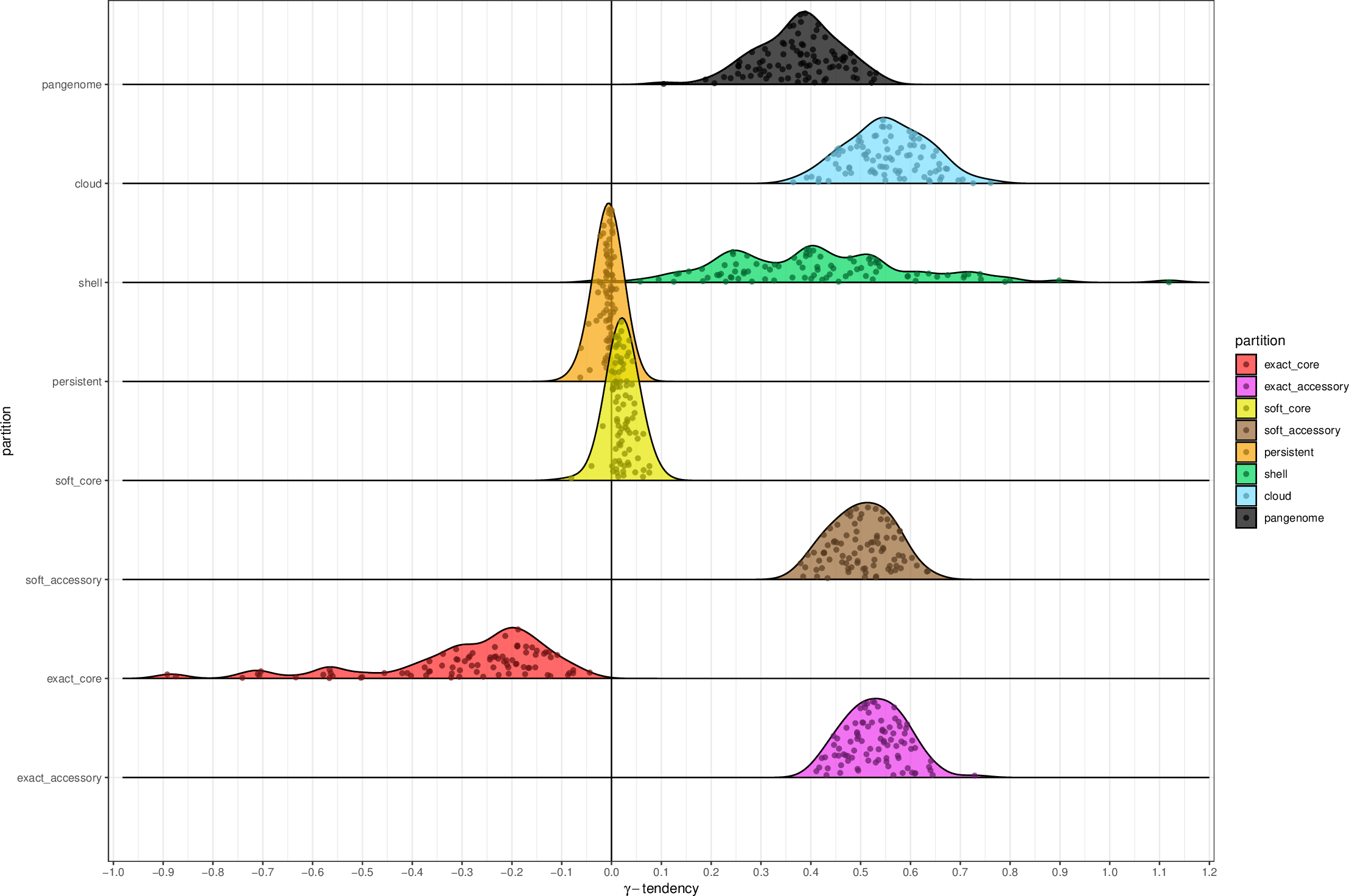

### S4 Fig

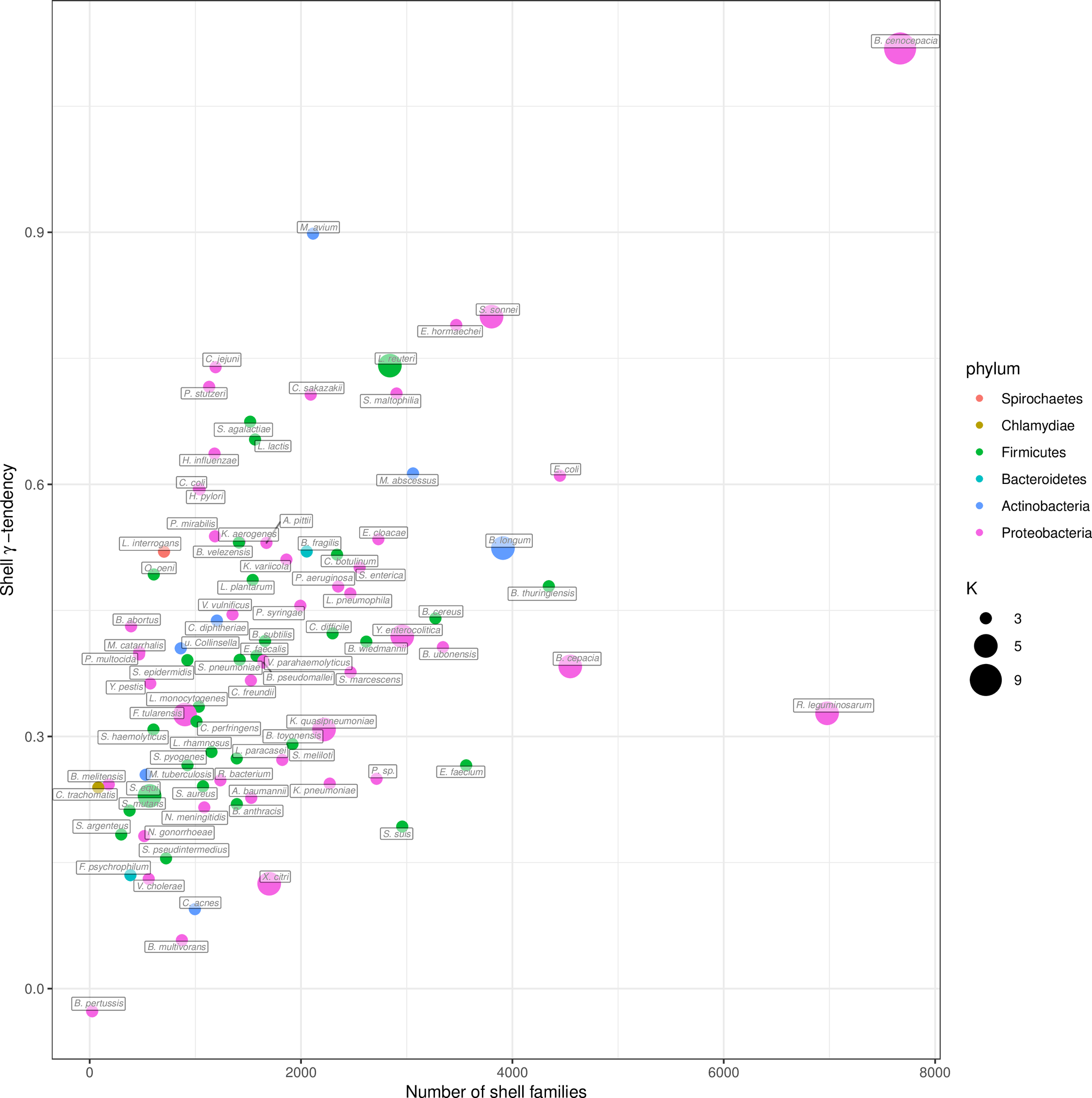

### S5 Fig

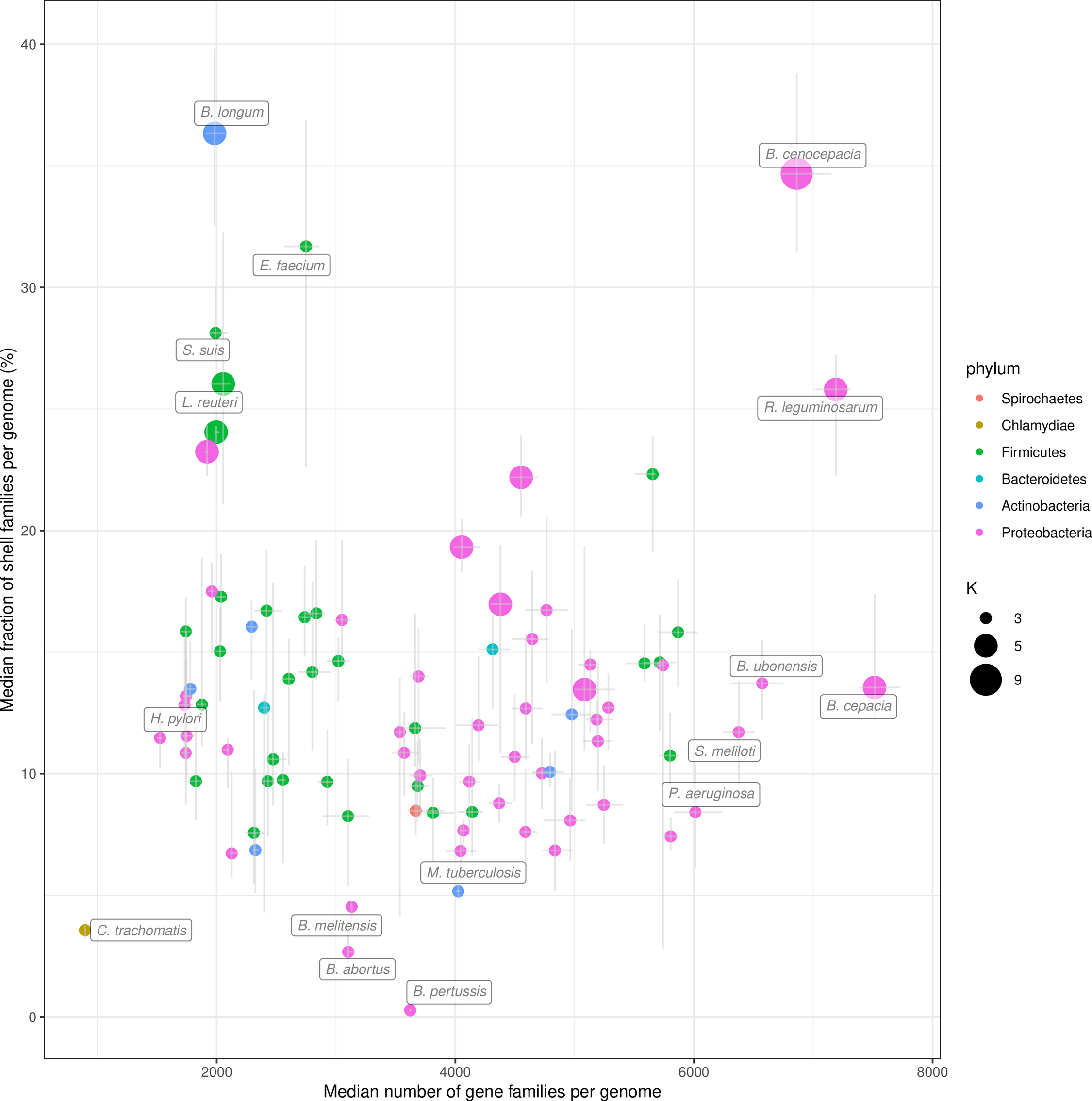

### S6 Fig

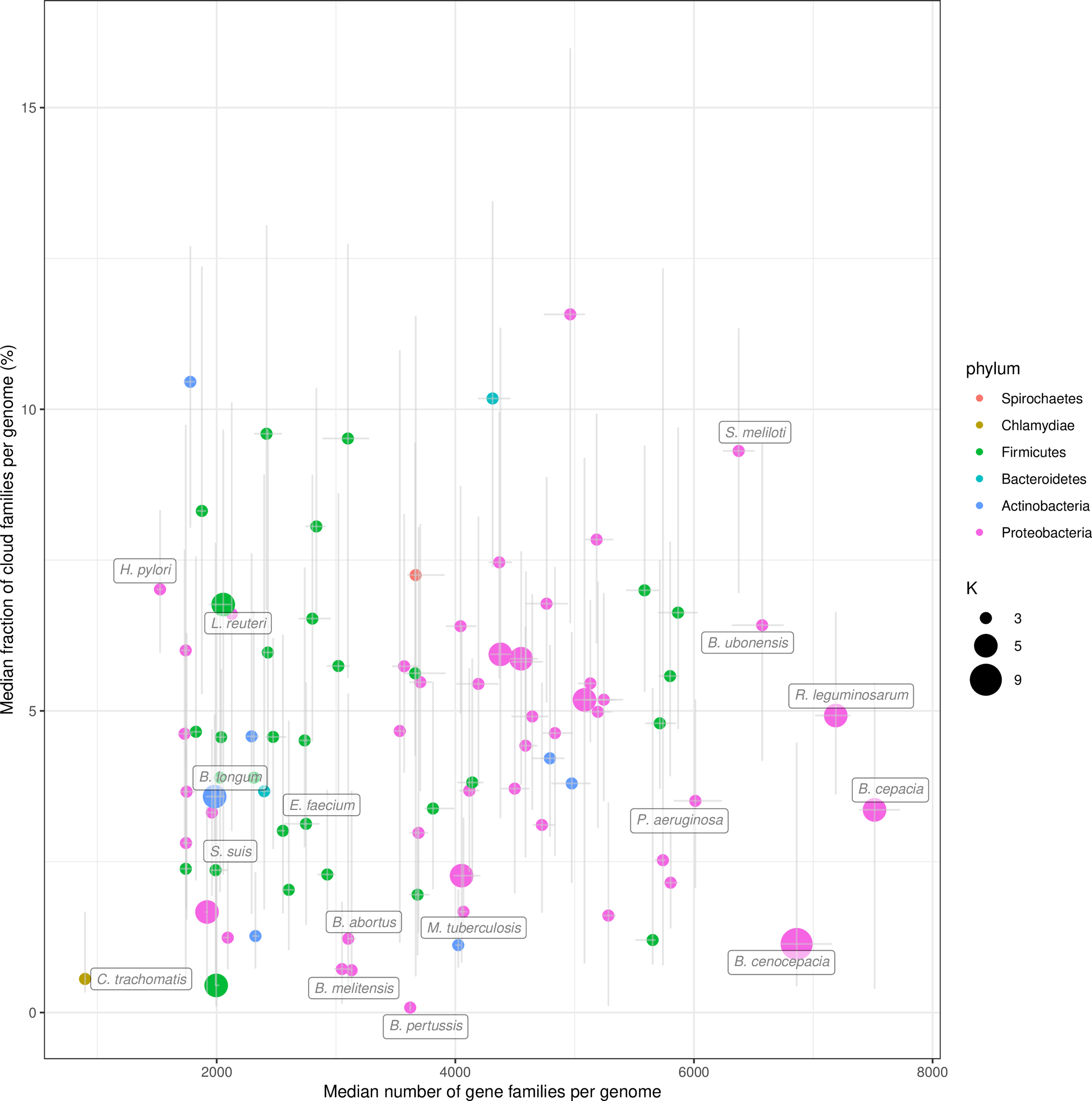

### S7 Fig

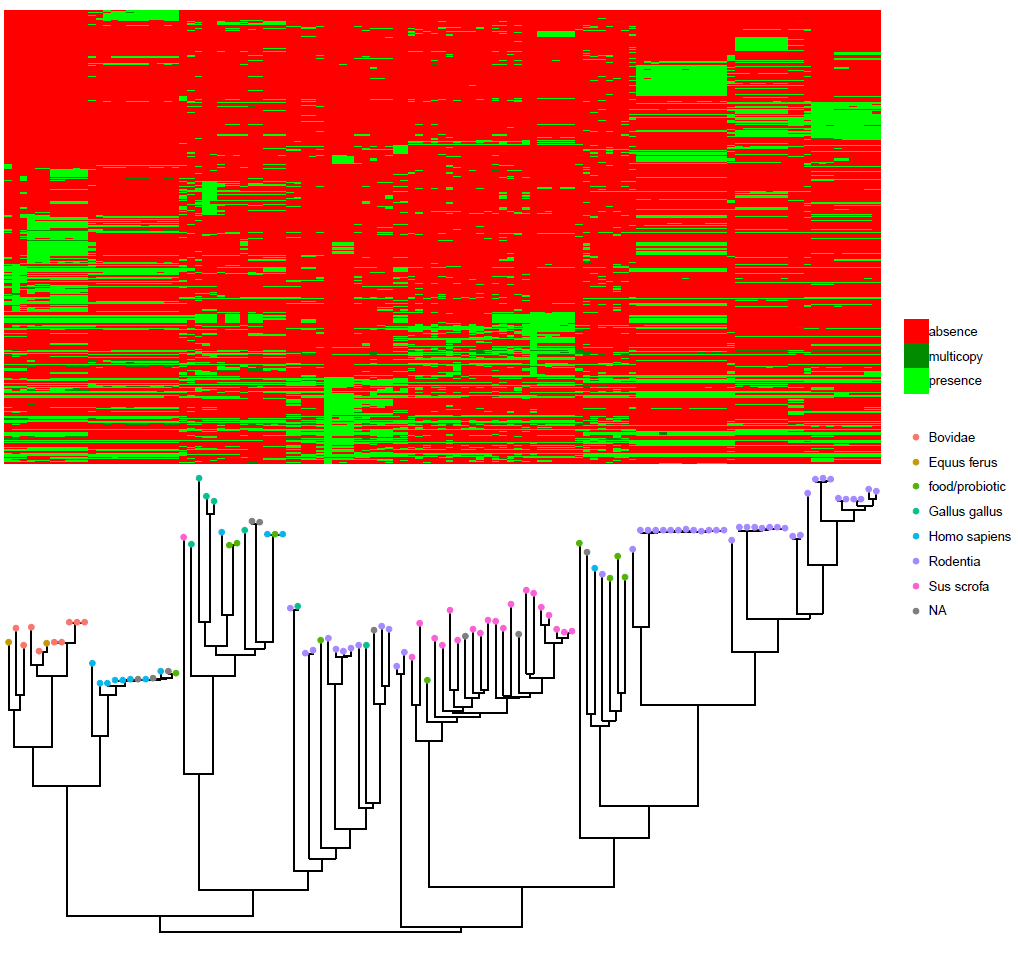

### S8 Fig

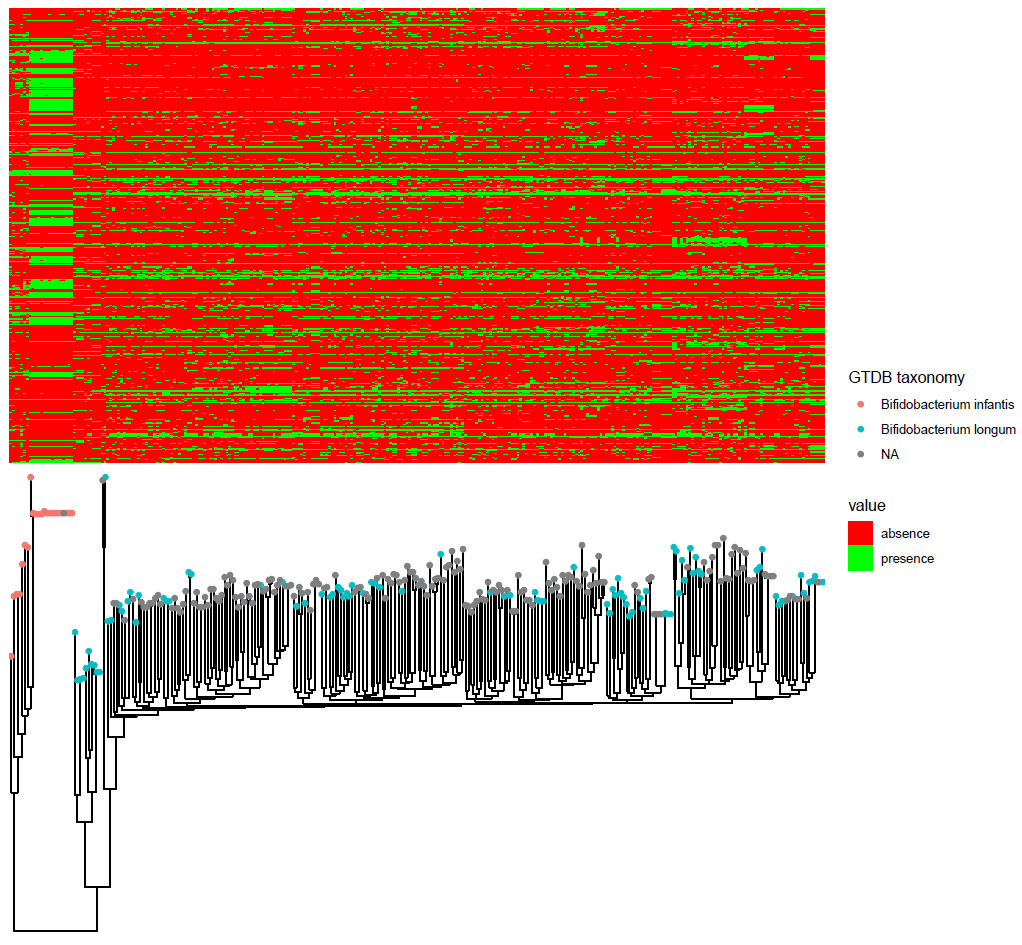

### S9 Fig

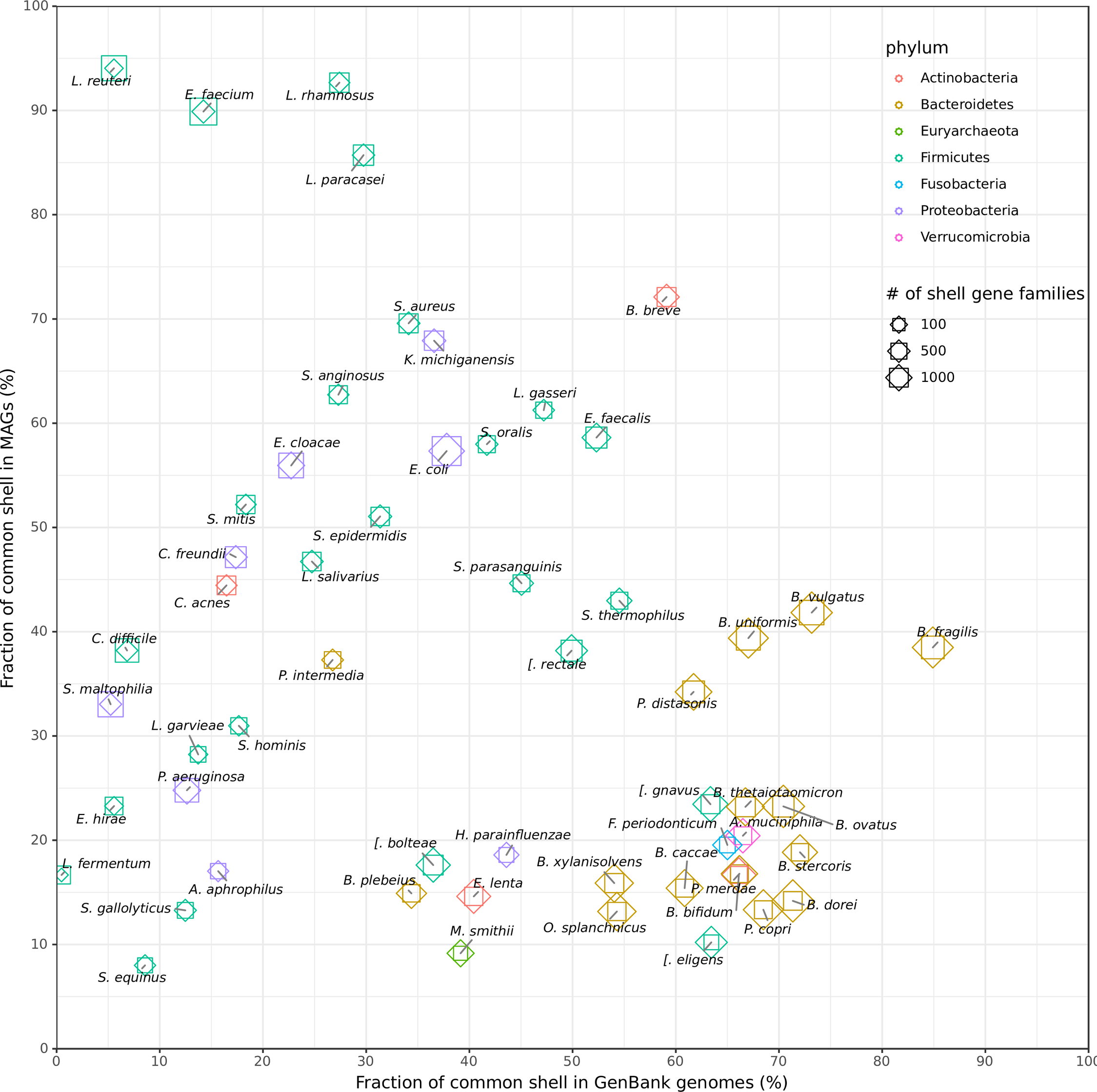
